# The Wooden Skull: An Innovation Through The Use Of Local Materials And Technology To Promote The Teaching And Learning Of Human Anatomy

**DOI:** 10.1101/2020.08.21.248476

**Authors:** Kintu Mugagga, Masilili G. Mwarisi, Samuel S. Dare

## Abstract

Skeleton models are important in facilitating student’s easy retention and recollection of information in the future. These may assist students carry out hands-on practice in order to acquire and practice new skills that are relevant to first aid.

The increasing number of medical institutions and medical students attracts the challenge of inadequate facilitation of the teaching and learning processes. This warrants a study and/or an exploration of an alternative solution such as wooden models in order to solve the problem of scarce and ethically restricted human teaching aids.

Wooden pieces (50cm *length* x 20cm *diameter*) from a ***Jacaranda mimosifolia*** tree were prepared for the carving process and wooden replicas of human skulls were made. Two experimental groups of randomly selected medical students (60: Active and 60: Control) were separately taught using wooden and natural skull models respectively. The two groups were assessed and evaluated using the natural skull models to compare their understanding of the Anatomy of the skull. Additionally, opinion statements were collected from participants in the active group during the oral examination.

Six (6) wooden skull models were produced and used for experimental study. Comparisons of academic scores (mean and median) between Active (students using the wooden skull) and control (students using natural skull) groups showed no statistical significant difference (P ≥ 0.05). Concerning the enhancement of learning skills; the wooden model was constructed in a way that would be able to enhance learning as it would be the natural skull.

The wooden skull model, with more improvement in structural formation, can adequately facilitate the teaching and learning of anatomy of the Human skull.

This project and the experimental study about utilization of the wooden skull model provides a good potential of using the wooden models to supplement the use of the natural human skull.

## INTRODUCTION

Human skeletal anatomy models, especially human skull anatomy models are great for patient education and students, in both educational and medical settings. Many classroom environments use skeletal systems to teach their students about the bone structure of the human body especially considering the complexity of structures in the skull. To understand difficult concepts, visual aids such as skeleton models are important in facilitating student’s easy retention and recollection of information in the future. Also, these skeleton models may assist students carry out hands-on practice in order to acquire and practice new skills that are relevant to first aid. In this regard, students in the early years of their medical training can practically demonstrate their ability for thorough and proper assessment to their teachers or examiners (Mentone educational, 2020).

The training of medical doctors in the preclinical years requires many dissections, but legal and ethical issues limits the availability of cadaveric material in many countries. Due to the scarcity of these skeleton materials, students have access to the materials for a limited duration. There is also the challenge of always transmitting unknown or known infectious agents from cadaveric skull (Mentone educational, 2020).

The use of models for teaching has been reported by several studies including the studying of muscular, cardiovascular, digestive systems and peripheral nerves by students who dissected cat which were observed to score lower marks than those who sculpted clay models of the body systems (Waters et al., 2005; DeHoff et al, 2011). Therefore it is important to note that the importance of anatomical models as learning tools in understanding complex three-dimensional (3D) anatomical structures cannot be overemphasized.

Over the past 30 years 3D printing has been used widely globally as described firstly by Charles W. Hull in 1986. 3D technology have been employed to construct several anatomical models such as bones, skull, heart, kidney etc. for educational purposes (Bizzotto et al., 2015; Narayanan et al., 2015; Knoedler et al., 2015; Costello et al., 2015). Although the skull has always remain the most complicated areas of anatomy, the use of skull base models in endoscopic training and temporal bone anatomy education for example is of great importance (Hochman et al., 2015). According to Chen et al., 2017, a randomized control trial (RCT) study designed to compare the learning effectiveness of 3D printed skulls with cadaveric skulls and the atlas revealed that study effectiveness especially the ability to recognize structures was enhanced better with the use of 3D printed skull compared with traditional learning materials.

The increasing number of medical institutions and medical professional students is practically a reality which positively address Medical Educational Partnership Initiative [MEPI]-Theme No.1, Increasing the Quantity and Quality of Health professionals (Mullan et al., 2012) and more globally supporting the MDGs (Kramer et al., 2008; Marieb & Mallet 2003). However, this quickly attracts major challenges particularly the facilitation of the teaching and learning processes. For any concerned teacher in any medical institution, effective and efficient teaching and learning is principally based on student centeredness, activity at classroom level and individualization amongst others (Harden RM and Laidlaw J.M, (2012). At present, a major challenge currently faced by Ugandan Medical institutions is the large class, often with over 300 biomedical students.

According to Sugand et al., 2010, the future of teaching medical students anatomy beyond the next ten years will probably be based largely on independent learning aids. With a ratio of 30 students/skull, this is 4 times less than the standard ratio of students per skull that is generally accepted (Malwadde et al., 2006). In terms of the costs, the price of the human skull is hard to define due to ethical values/restrictions. The hardship involved in acquiring the human model and the ethical rules governing its utilization outweighs any form of pricing system and thus makes it hard to attach the real monetary value. Although plastic models have also been adapted but the cost of acquiring them is enormous for institutions in a developing country like Uganda. Nevertheless, there is a consistent increase in the interest to develop and employ new educational tools for teaching since it has been established that the use of visual aids like models enables students to perform better in anatomy course examination (McNutty et al., 2010).

Comparatively, the wooden model once accurately constructed, the carvists and the anatomists using their mastery, can generate as many models as required at low cost as $100 per wooden skull. With about 12 human skulls available at gross anatomy laboratory at African institutions, this leads to student /skull ratio of 30 which is far less the standard ratio accepted. Thus, this warrants a study and an exploration of an alternative solution to the problem of scarce and ethically restricted human teaching aids.

*J*. *mimosifolia* is a spectacular tree found in many tropical and sub-tropical countries. Similar to many other ornamental trees, it is regarded as native to South America however, it has spread widely over the century, naturalized in many countries and also penetrated into many locations in East Africa countries including Uganda (Witt and Luke 2017; GBIF 2014; Dharani, 2005). *J*. *mimosifolia* is an invaluable tree having impact economically, socially and environmentally as such, it is useful as carvings and tool handles, interior carpentry wood source for fences and fuel plant Dharani, 2005). In this regard, we chose *J*. *mimosifolia* because the timber is yellowish-white, moderately heavy but hard and easy to work with.

## MATERIALS AND METHODS

### The Wood source: *Jacaranda Mimosefolia*

Building our skull model begins with a wood source, *Jacaranda Mimosefolia* (Figure 1) which was harvested from Nyendo village, Masaka district, Western Uganda. Our choice was based on its physical properties namely: the yellowish-white color, moderately heavy, fine textured, attractive, close grained and figured. Additionally *J*. *mimosifolia* has good working qualities like: lightness, easy to work with, it stains, glues and sands easily. Its density is established to be 0.615 g/CC (Witt and Luke 2017; GBIF 2014; Dharani, 2005).

**Figure 1:**
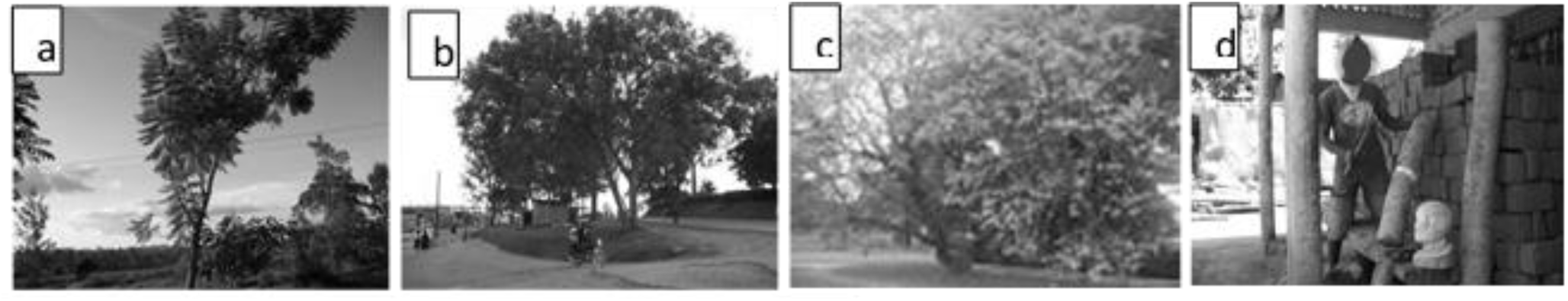
The Wood Source and Cutting into Logs

### The Wood Processing

We cut with a hand saw 10 cylindrical pieces (50 cm in length and 20cm in diameter) each from a 10m Jacaranda tree [mature height]. We dried the pieces under shade and debarked them afterwards using Stanley chisels (12-25mm). After weight reduction, down to approximately 9.5 Kgs in 30 days, we subjected them to primary coarse-carving (Figure 1 & 2A). We carried out the wood processing at the department of human anatomy, Kampala International University, Ishaka, Campus in collaborations with wood carvists of Kampi Art And Crafts Wood Carving Center Kitovu, Masaka Uganda.

### The Carving Tools

We were guided by the wood carvists and an assortment of carving tool were bought from different sources in Kampala. However, a good number of them could not be found on the market. Collaboratively, we hired the workshop tools from Kampi carvists under a contractual agreement. The following tools were used: Beginner carving tool set, wood carving kit, whittling knife kit, whittling jack, ultimate power sharpener, palm carving tools, micro carving tools, miniature carver, power chisel, angle grinders, brick and mortar saw, round and oval eye punches, carving scraper set, mallet size set, digital calipers and others (Figure 2D, E & F).

**Figure 2:**
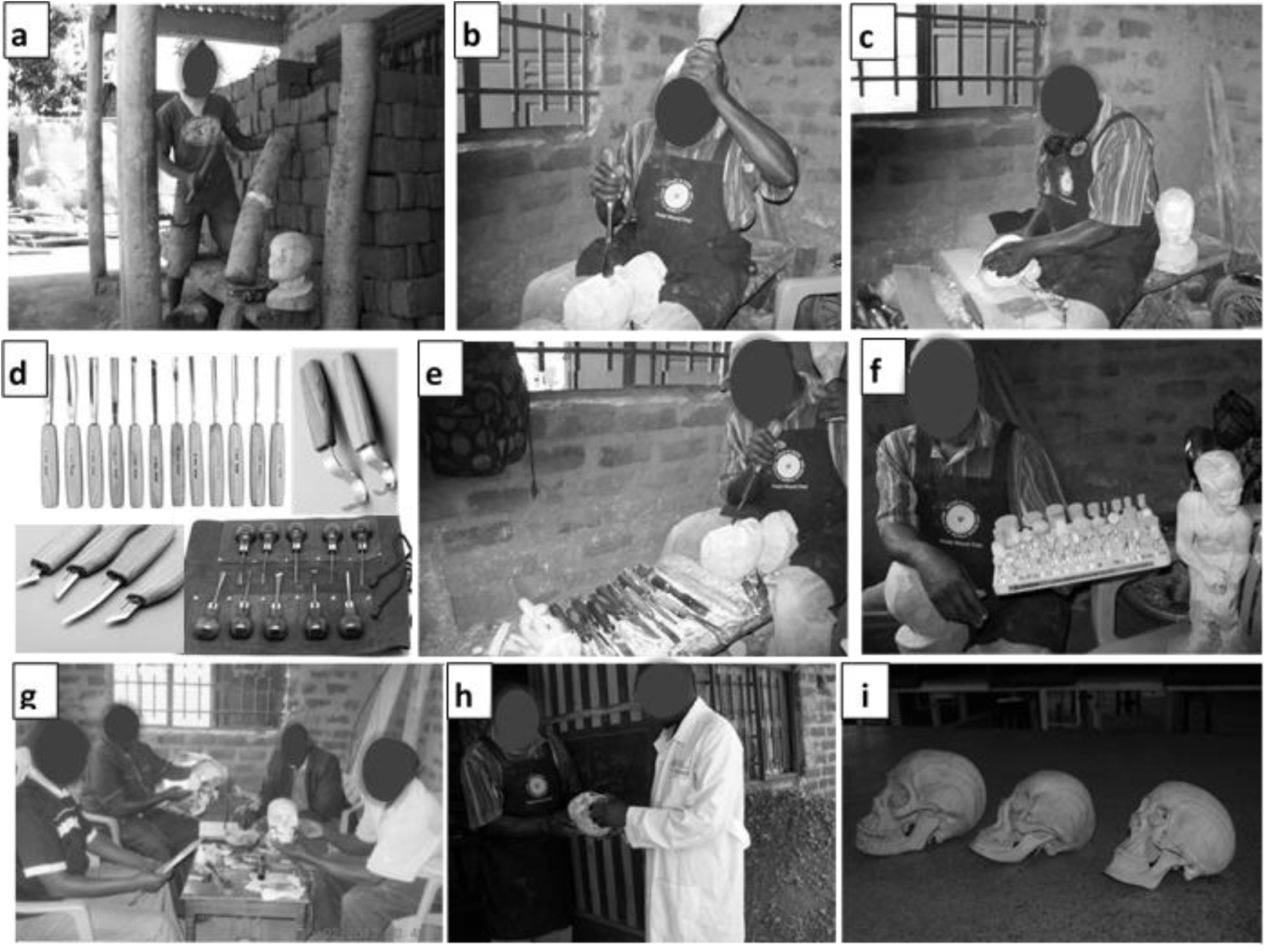
Wood Carving Process, Tools, Technical Team and Finished Product

### The Carving Process

The proportionate dimensions of the wooden skull model under transformation was based on the anatomical guidelines provided by the Medical illustrator (*Mr*. *Paul Lukiza-Makerere College of Health Sciences*) (Figure 2B, C, E & H).

### The Guiding Models

For perfection, we used the following reference models: i) An exploded human skull model, ii) an intact (complete human skull), iii) a hemi (sagittal) section of the human skull, iv) a human skull without skull cap (Calvaria) exposing the cranial cavity and v) Human Anatomy Text books (Marieb, 2003), showing the elaborate Anatomy of the human skull (Figure 2G).

### The Carving output

After, 8 weeks (including 4 weeks of seasoning period) we successfully produced six (6) wooden Skull models (Figure 2H & I). On average each skull took 1week of carving. The carved skull models were used for the experimental study with the biomedical science students.

### Experimental Study to validate the use of Wooden Skull for Anatomy Education

We randomly selected two groups of medical students (60 in each group), from the MBChB, Class (Year 2-semester 1). The two groups were fully notified about the details and intentions of the experimental study. They were briefed about their rights as participants in the study and their consent and willingness were sought for according to required guidelines.

We taught the anatomy of the human skull to the two groups separately using wooden and the natural models for the active and control groups respectively in using gross anatomy laboratory 1 and gross anatomy Laboratory 2 respectively. The teaching took place in May, 2013 for 24hrs in 4 days covering Skull development, Splanchnocranium, Neurocranium, Scalp, muscles and Joints. The periods covered theory and practical concerns. The two groups were equally facilitated by two anatomists i.e each group had two anatomy lecturers [ratio 30 students/lecturer] who used similar curricular guidelines to cover the topics. The student /skull ratio was for both groups 10: 1 and every lecture was followed by a practical session. Due to limitations in time, the carving of the cranial cavity was omitted and concentration of the study and assessment was limited to external anatomical details on the wooden skull model.

### Assessment

We comparatively assessed the two groups by subjecting them to a standard exam which had written, practical and oral sections. The examination was based on the traditional natural human skull model and proportionally, the practical scored 50%, written 30% and oral 20%. Additionally, an opinion statement about the wooden skull [as compared to the natural skull] was collected from participants in the active group during the oral examination.

### Statistical Analysis

For data analysis using SPSS (version 18), comparative scores of the two groups (from the written, practical and oral exams) were analyzed using two-sample t test with equal variances. The chi-square test was used to determine the statistical difference between the two sets of scores. Finally, comparative scores between the two groups showed no significant difference (P ≥ 0.05).

A descriptive statistics were employed to broadly evaluate the effect of treatment group on student perceptions about the activity and science. Student’s open-ended responses were qualitatively analyzed using an inductive reasoning where related responses were grouped into subsets that are quantifiable. In this respect, the responses were categorized by the researchers independently and reached agreement of not less than 90% on their categories after further discussion (Maykut and Morehouse, 1994).

### Ethical Considerations

The study was approved by Mbarara University of Science and Technology [MUST] **Institutional Review Board** IRB [No. 08/09-12] and Recommended for registration with Uganda National Council for Science and Technology.

## RESULTS

### Appearance of Cranial Features

The appearance of our wooden skull specimen model demonstrated the anatomical features of the natural skull (figure 3). Most of the bones and sutures such as coronal, squamosal, lambdoid, parietomastoid were clearly shown. Other features clearly demonstrated are orbital cavity, nasal cavity, pterion, mastoid process, mandible and upper & lower jaws, zygomatic arch, supra and infratemporal fossa, temporal lines temporomandibular joint, occipital protuberances etc.

**Figure 3:**
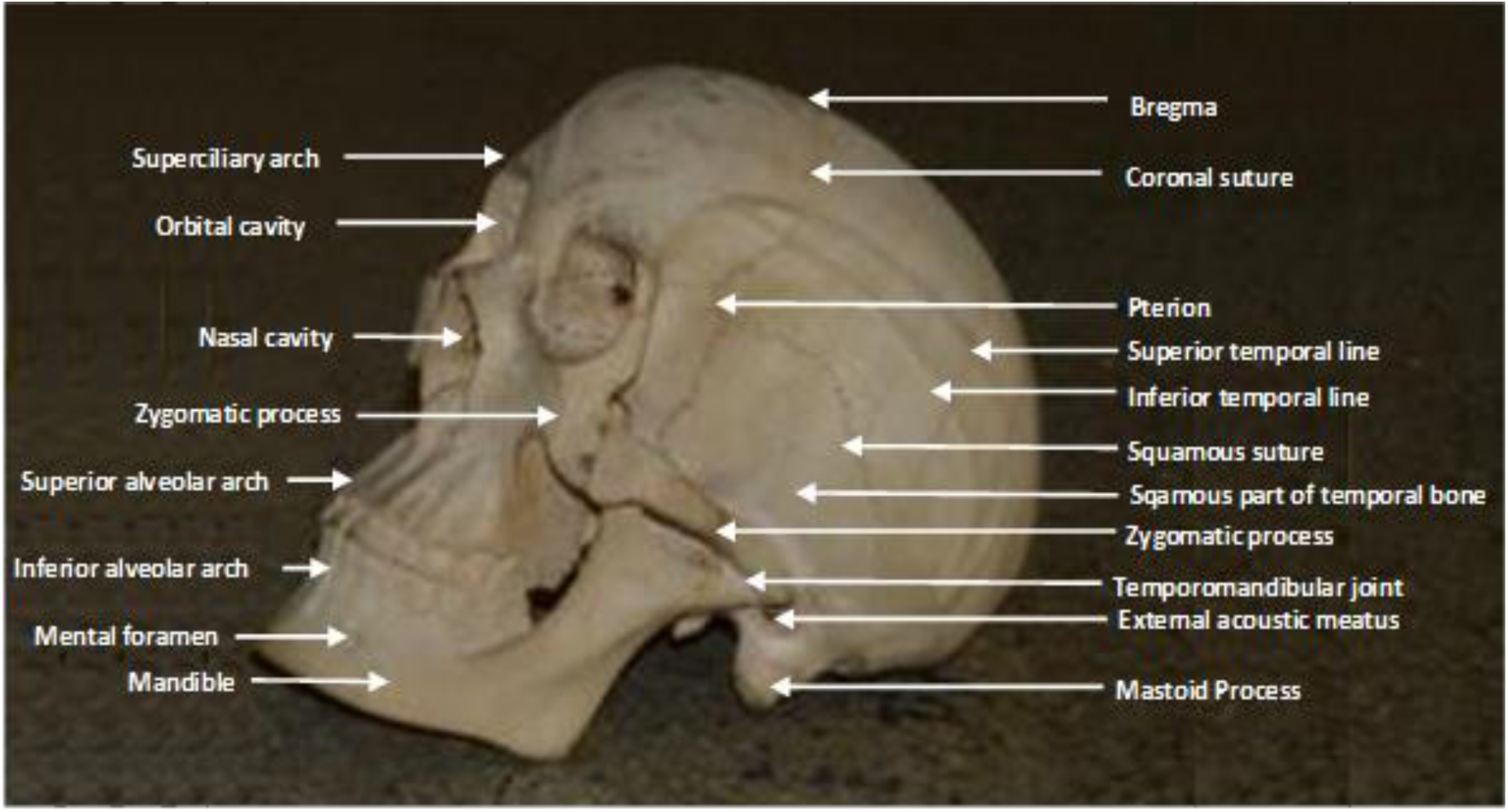
Our wooden skull with some anatomical features labelled

### Null hypothesis

Teaching the anatomy of the human skull by using an artificial model provides similar exposure and understanding as the cadaveric model. To test the hypothesis, we compared the group exposed to human skull with the one exposed to wooden skull, using scores from written, oral and practical examination. We set the level of significance 0.05.

### Students’ Quantitative Evaluation

To understand the failure rate of the students, we categorized the scores as 40-50 (failed), 51-60 (pass), 61-70 (credit), 71-80 (B), and 81-100 (A). We used chi square to determine the difference.

## Students’ Qualitative Evaluation

### Learners Reaction, Knowledge Acquisition, Skills, Attitudes and Behavioural Changes through converted oral exam transcripts

#### Qualitative findings

The innovation of a wooden skull model has been presented in 2 perspectives:

A: Assessment of the anatomical features of the wooden skull

B: Comparison of wooden skull with the natural skull

##### A: Assessment of the anatomical features of the wooden skull

**Theme 1: Fixed joints**

To innovate a model for anatomical learning experience from wood, takes great courage, skills and knowledge. Framing a skull from wood can equally bring out a structure like that of a natural skull. Though some structures are not easy to carve out exactly as the natural skull therefore some parts like the joints may be fixed at certain places. According to respondents:

> *“…The wooden skull has a fixed temporomandibular joint*.*”*

**Theme 2: Clear demonstration of sutures**

The sutures were clearly carved out, may be due to their anatomical make up which made them be easy to carve from wood. This made them be clearly identified by the respondents during their learning experience using a wooden skull. According to respondents:

> *“*… *Lambdoidal suture and sagittal sutures are clearly demonstrated*.*”*

**Theme 3: Enhancement of learning skills**.

The wooden model was constructed in a way that would be able to enhance learning as it would be the natural skull. One of the respondents stated:

> *“Personally I have enjoyed learning using this model* … *it has increased my desire to correlate the structures and be able even to know more concerning the true skull*.*”*

##### B: Comparison of wooden skull to natural skull

**Theme 4: Close connection**

The wooden skull was constructed to have similar structures as the natural skull. This created a close connection and similarity to the natural skull. A respondent revealed that;

> *“There is close connection between the artificial model and the wooden model*.*”*

**Theme 5: Easy to use for learning**

Compared to the natural skull, the wooden skull was easy to use in learning. This was due to the simplicity of how it was carved. In this regard, respondents stated;

> *“*…*one thing it has caused, is my attraction to the structures of the skull and been able to increase my desire to know more about the skull”*. Also
>
> *“I find the skull model easy to study and learn*…*”*

## DISCUSSION

Supported by Harden & Laidlaw (2012) observation in clinical training and assessment where multiple difficulties are encountered in standardizing real patients thus leading to development of simulated patients, the use of natural/ human models in quality teaching of human anatomy is characterized by limited choice for: i) the best model ii) how many models can be acquired as needed or iii) how best to standardize the existing models.

The output of this experimental study about utilization of the wooden skull model clearly signifies the potential of using the wooden models to largely substitute the natural human skull. This is because examination results of our study showed no statistical significant difference between the groups of students exposed to the use of both the natural skull and the wooden skull model (Table 1 – 4). Considering the mark distribution of the students’ performance, there was no significance difference between the control and active groups in terms of pass and failure rate (Table 5 & 6).

**Table 1:** Written Exam. Two-sample t test with equal variances

| Group | Obs | Mean | Std. Err. | Std. Dev. | [95% Conf. Interval] |  |
| --- | --- | --- | --- | --- | --- | --- |
| Active | 50 | 21.93 | .5350968 | 3.783706 | 20.85468 | 23.00532 |
| Control | 42 | 21.88095 | .5270856 | 3.415905 | 20.81648 | 22.94542 |
| Combined | 92 | 21.90761 | .3754222 | 3.600923 | 21.16188 | 22.65334 |
| diff |  | .0490476 | .7578572 | -1.456568 | 1.554663 |  |

P = 0.53, t = 0.0647, degrees of freedom = 90
Based on the above p value of 0.53, there is no significant difference in performance of students exposed to human skull with the ones exposed to wooden skull in a written exam. Note that the confidence intervals also overlap while the mean is also the same.

**Table 2:** Practical Exam. Two-sample t test with equal variances

| Group | Obs | Mean | Std. Err. | Std. Dev. | [95% Conf. Interval] |  |
| --- | --- | --- | --- | --- | --- | --- |
| Active | 50 | 36.65 | .8742635 | 6.181977 | 34.8931 | 38.4069 |
| Control | 42 | 36.52381 | .8921116 | 5.781544 | 34.72215 | 38.32547 |
| Combined | 92 | 36.59239 | .6224292 | 5.970131 | 35.35601 | 37.82877 |
| diff |  | .1261905 | 1.256444 | -2.369956 | 2.622336 |  |

P = 0.54, t = 0.1004, degrees of freedom = 90
Based on the above p value of 0.54, there is no significant difference in performance of students exposed to human skull with the ones exposed to wooden skull in a practical exam. Note that the confidence intervals also overlap while the mean is also the same.

**Table 3:** Oral Exam. Two-sample t test with equal variances

| Group | Obs | Mean | Std. Err. | Std. Dev. | [95% Conf. Interval] |  |
| --- | --- | --- | --- | --- | --- | --- |
| Active | 50 | 14.66 | .35209 | 2.489652 | 13.95245 | 15.36755 |
| Control | 42 | 14.61905 | .3507953 | 2.273413 | 13.9106 | 15.32749 |
| Combined | 92 | 14.6413 | .2481809 | 2.380468 | 14.14832 | 15.13429 |
| diff |  | .0409524 | .500991 | -.9543536 | 1.036258 |  |

P = 0.53, t = 0.0817, degrees of freedom = 90
Based on the above p value of 0.53, there is no significant difference in performance of students exposed to human skull with the ones exposed to wooden skull in an oral exam. Note that the confidence intervals also overlap while the mean is also the same.

**Table 4:** Total Aggregate Score. Two-sample t test with equal variances

| Group | Obs | Mean | Std. Err. | Std. Dev. | [95% Conf. Interval] |  |
| --- | --- | --- | --- | --- | --- | --- |
| Active | 50 | 73.32 | 1.746937 | 12.35271 | 69.8094 | 76.8306 |
| Control | 42 | 73.02381 | 1.765421 | 11.44123 | 69.45847 | 76.58915 |
| Combined | 92 | 73.18478 | 1.238702 | 11.88122 | 70.72425 | 75.64531 |
| diff |  | .2961905 | 2.500407 | -4.671305 | 5.263686 |  |

P = 0.55, t = 0.1185, degrees of freedom = 90
Based on the above p value of 0.55, there is no significant difference in performance of students exposed to human skull with the ones exposed to wooden skull in the overall exam. Note that the confidence intervals also overlap while the mean is also the same.

**Table 5.**
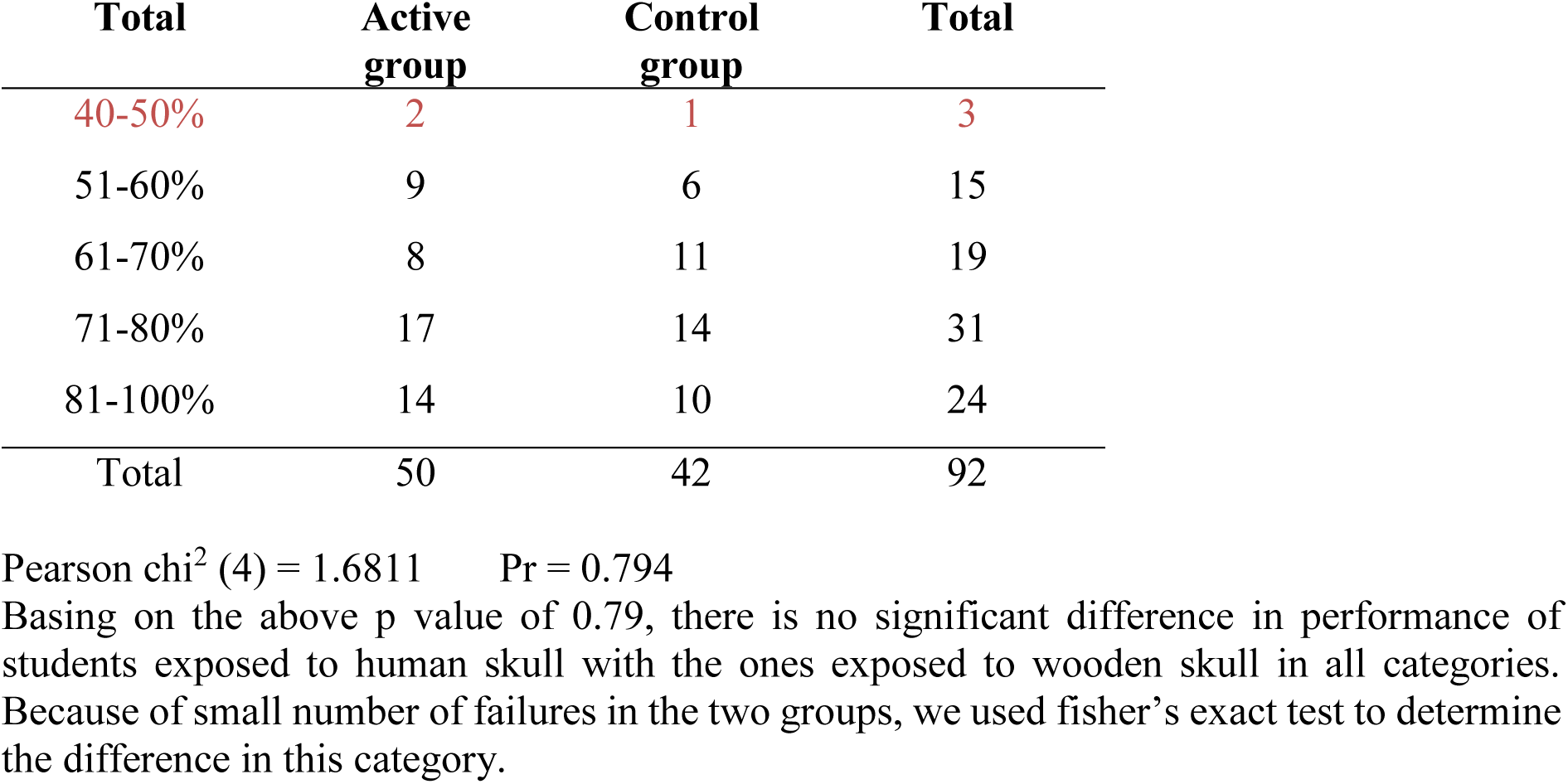

**Table 6.**
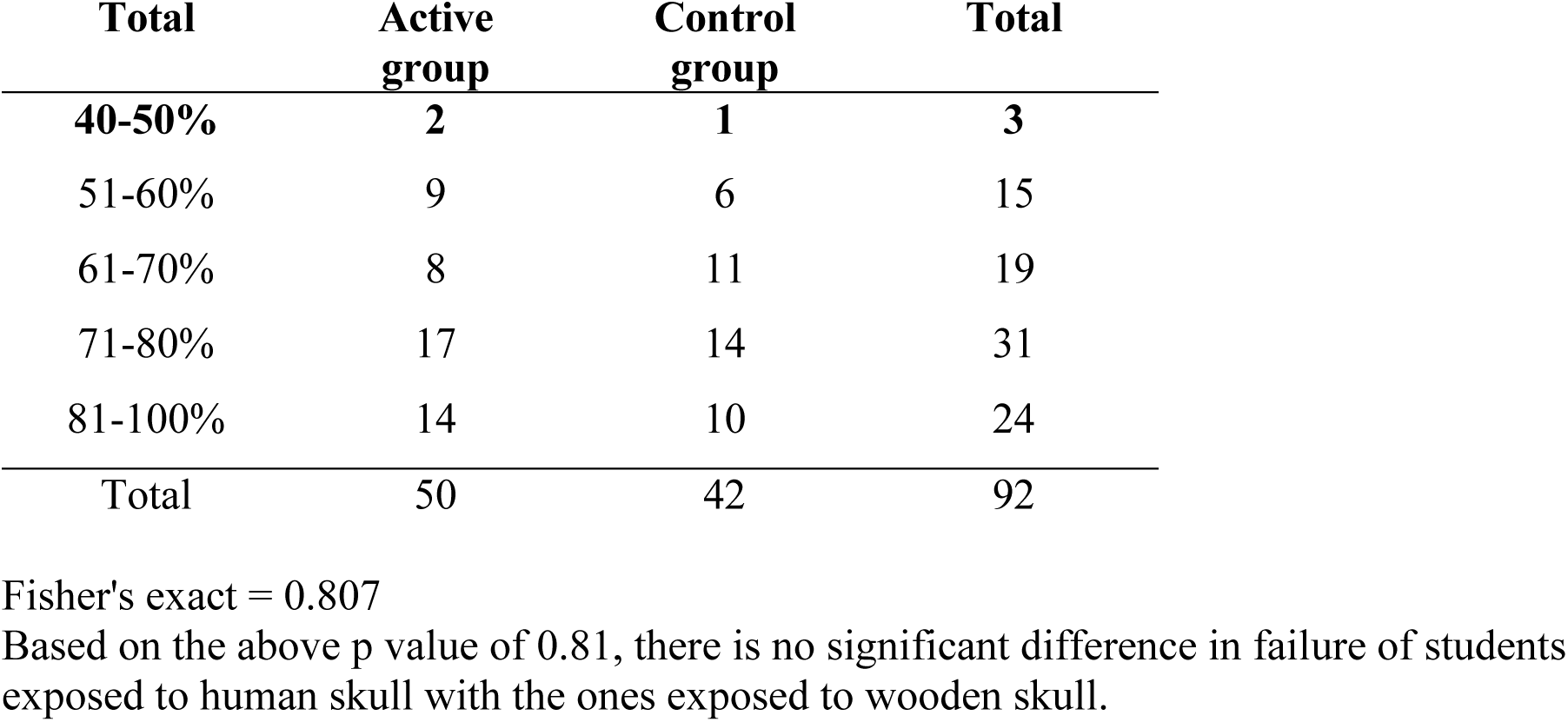

According to Ruiz et al., 2006, the use of physical learning objects are necessary for robust learning in both preclinical and clinical education and as well furnishes several research opportunities. Embracing learning objects for adaptive learning can easily help to measure the reaction of learners, knowledge acquisition, skills and attitude as well as behavioural changes. Therefore, transcribed the oral exam of the students as shown in section B of our result revealed no significant difference between the Cadaver skull and the wooden skull.

Our observation agreed with the publication of Andreas Vesalius in 1543, “De humani corporis fabrica” which brought a rebirth in anatomy in Europe as reported by Russell in 1972. This publication highlighted the need for some representation of the body anatomy in the form of image picture and models due to lack of preservative means and scarcity of bodies for dissection. Though the need was partially met by anatomical prints, nevertheless it was not satisfactory because a single-dimensional picture cannot give the true impression of the body structure as required to for an untrained person. Obviously a three-dimensional figure will be of advantage to a single-dimensional picture despite that it may not reveal details as perfect as it would in the engraving (Russell 1972).

Khot et al., 2013 in an experimental study, compared the results of a knowledge test between learners who studied female pelvic anatomy using a solid 3D plastic model, a static atlas-type compendium of photographic images and computer-based virtual reality materials. The results revealed that the group of students that used the plastic model remarkably performed better than the other groups which used photographic images and the computer based virtual materials. Also Daniel Preece and his mentors Drs. Sarah Williams, Richard Lamb, and Renate Weller in their article reported that students who used plastic model of an equine foot learnt extremely better than those who used textbooks or 3D computer models (Preece et al., 2013).

Our results shows wooden skull model with the features demonstrated (figure 3) and been user friendly presented a significant impact on the student’s attitudes, learning and retention of the osteology of the skull without statistical significant difference in the mark distribution between the control and the active groups (figure 4 and 5). It is also important to note that the psychological disorder or problems that the natural skull might bring to the students causing them to withdraw from its use and learning well can be solved with the use of the wooden skull model therefore better satisfying the requirement for teaching and learning.

**Figure 4:**
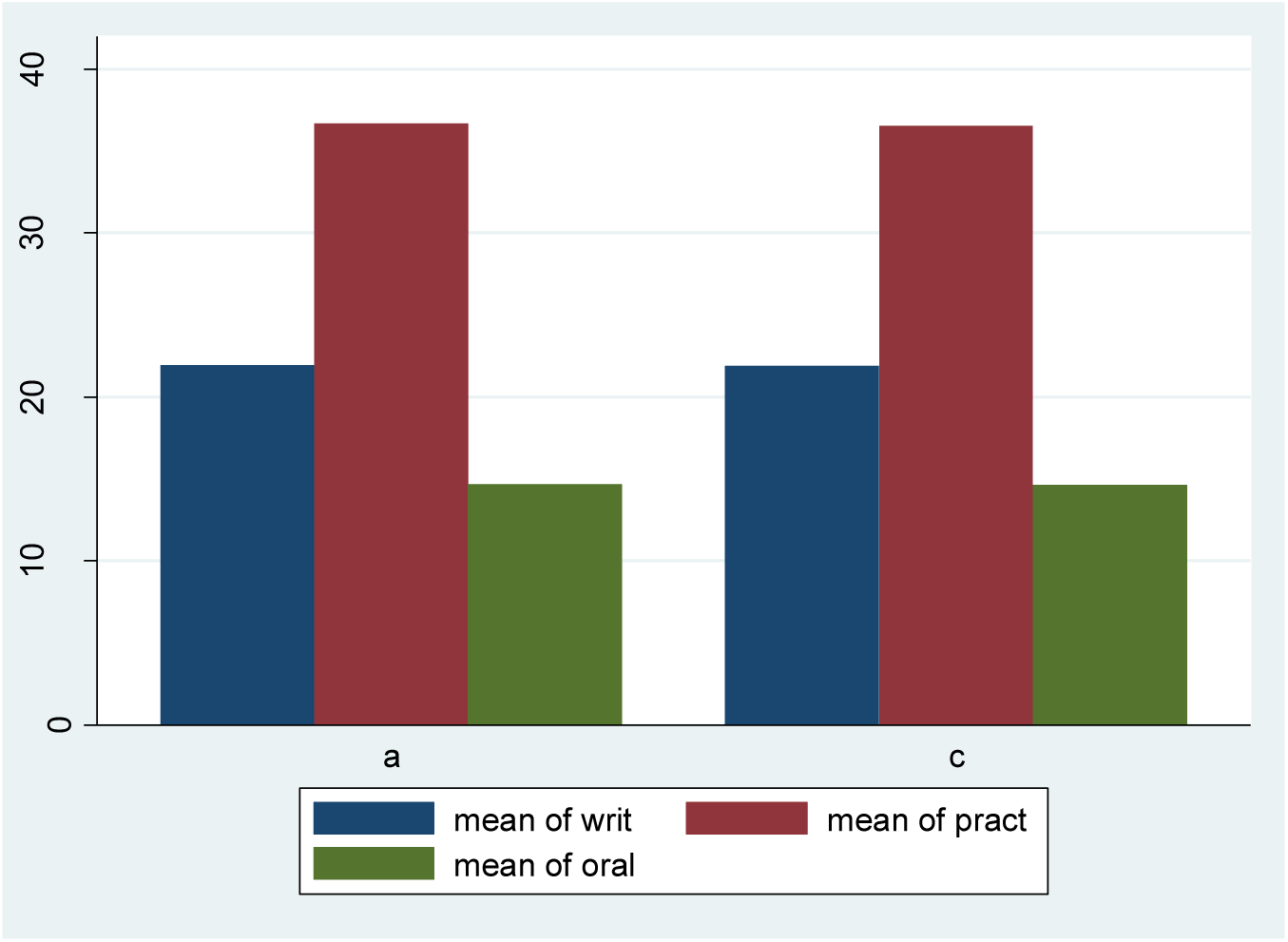
Mean comparison of Active (a) and Control (c) groups in academic performance Bar chart showing mean values of student test scores for the written, practical and oral exams for each treatment (natural skull and wooden skull)

**Figure 5:**
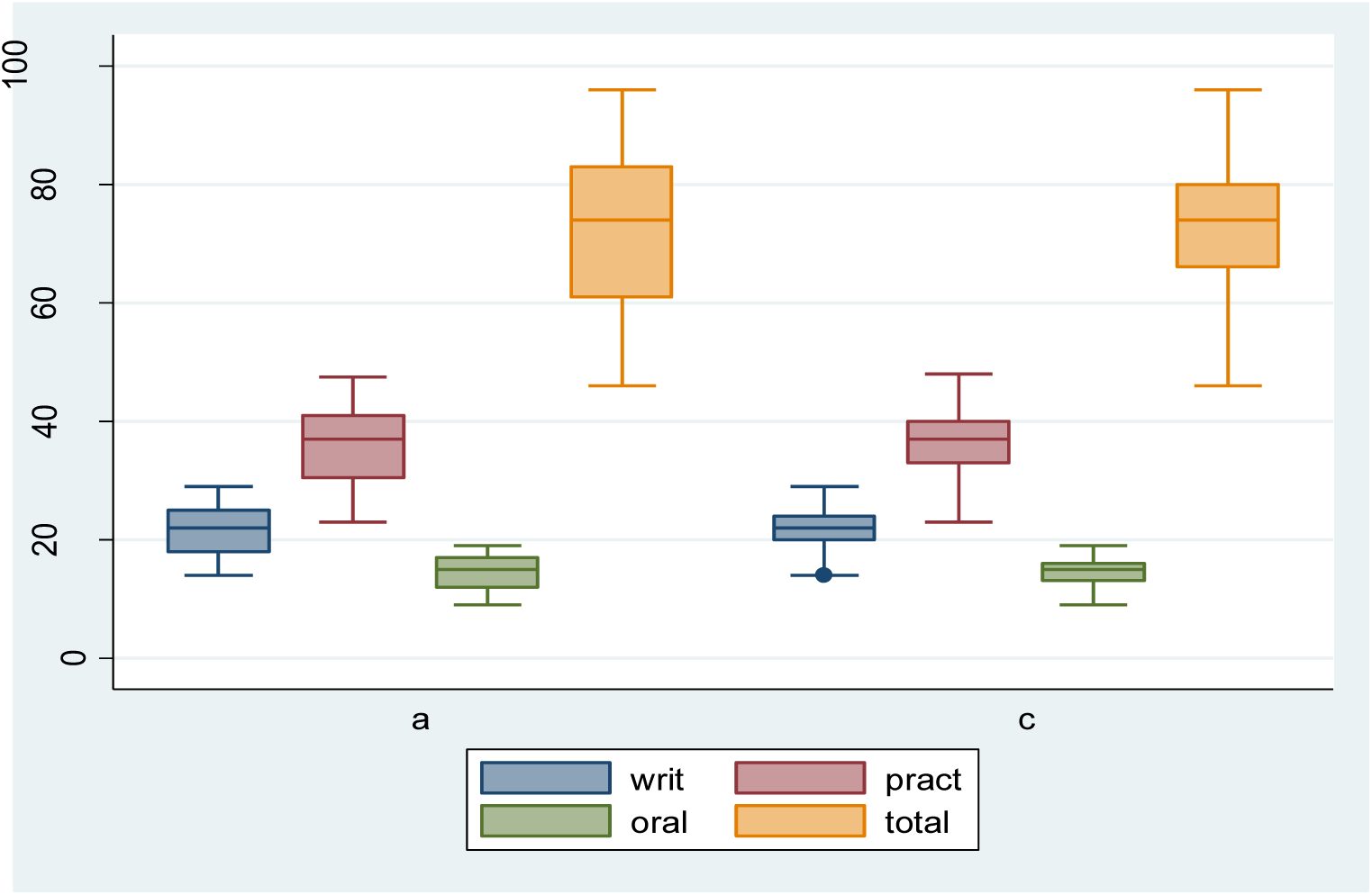
Median comparison of Active (a) and Control (c) groups in academic performance Box plot illustrating the median, minimum, and maximum as well as 25–75 percentile ranges of student test scores for the written, practical and oral exams for each treatment (natural skull and wooden skull).

### Practical Significance

This wooden skull model shall serve as starting point in setting up the human anatomy skills lab, which shall improve on availability of teaching models for medical students teaching and learning. With this production, at least one skull model for every 5 trainees can be achieved; i.e even a class of 300 learners can ably have 60 skull models available for training. Desired quality teaching and learning practices like: Small groups, student centeredness, individualization, activity etc, shall be easily achieved.

### Sustainability

#### Growing expertise

We have demonstrable evidence that the team that practically participated in the carving process (the wood carvists, medical illustrator and the anatomists) is now more versed with knowledge and art of fabricating and teaching the anatomy of the skull.

#### Source of raw material

Wood being a renewable resource, its continued availability is about growing the tree of choice (*J*. *mimosifolia* tree) which truly comes along with many other benefits.

#### Durability of the Wooden Model

Jacaranda wood has a property of durability based on its physical and chemical composition. The models are still treatable with ordinary wood preservative (*like Wood-bliss*) to prevent microbial (fungal, insect etc) degradation under the laboratory storage and usage environment.

#### Environmental friendliness (Compatibility with MDG-7)

We cannot ignore the contribution to the green environment as we grow more jacaranda trees as future source of wood for the models (Sachs & McArthur, 2005). This is further supported by the high conversion rate (every 10m tree produces twenty 50cm pieces, where each piece is convertible into one wooden skull model i.e 20 models per tree). More importantly, wood carving process is purely mechanical with no chemical involvement (i.e no addition or emissions of chemical or gases to the environment). Contrary to Gregory (2009) which highlighted that plastics are manufactures with chemicals that are known to be toxic. Studies in animal model organisms reported that that chemicals used in the manufacture of plastics and present in human population have potential adverse health effects (Talsness et al. 2009). These chemicals burden correlates with adverse health effects such as reproductive abnormalities in human population (Swan et al. 2005; Swan 2008).

As one of the most naturally renewable energy sources known, wood has less impact on the environment than other materials. It can also last longer than a lifetime when treated correctly and waste from production of wooden materials is limited and one hundred percent degradable. Given that wooden pallets are more environmentally friendly and more eco-friendly than plastic pallets due to current concerns about climate change (Sebastian et al., 2020; Penn State 2020), this wooden skull model carved from *J*. *mimiosifolia*, though not easy or simple to illustrate all the structures clearly especially the internal structures, may be a better alternative to plastic models. The researcher, therefore look forward to overcoming the limitations encountered in the process of carving subsequently by ensuring to improve on the external features as well as properly carving out the internal structures to effectively demonstrate in details the anatomical structures with minimal defects.

## CONCLUSION

The outcome of this project and the experimental study about utilization of the wooden skull model at Kampala International University, Anatomy Department, provides a good potential of using the wooden models to supplement the use of the natural human skull. The use of the wooden model for training medical students can be done anywhere and anytime without probable exposure to infection from the cadaver skull and also freedom from ethical and legal restrictions.

## DATA AVAILABILITY

The metadata used to support the findings of this study are available from the corresponding author upon request.

## ACKNOWLEDGEMENTS

The authors gratefully acknowledge Prof. Jonathan Nshaho, the MEPI/MESAU Co-PI at Kampala International University, Prof. Nelson K. Sewankambo – MESAU PI), the students who participated in the experimental study and the rest of the staff of anatomy department who ensured a conducive environment. We also appreciate the Annals of Global Health which had published our abstract with the title of this article from a poster presentation at the 5^th^ Annual Consortium of Universities for Global Health (CUGH) Conference (2014) in Washington, DC.

